# Environmentally-determined symbiont communities highlight flexibility of Aiptasia-algal symbiosis

**DOI:** 10.64898/2026.05.11.724104

**Authors:** Maria Ruggeri, Connie S. Machuca, Samuel A. Bedgood, Stacy A. Krueger-Hadfield, Carly D. Kenkel

## Abstract

The mechanisms driving host-symbiont associations across space and time in contemporary mutualisms can give insight into the capacity for symbiotic organisms to respond to environmental change. High specificity between partners can increase cooperation and facilitate efficient holobiont selection, whereas low specificity could reduce host benefit, but facilitate adaptive associations across heterogeneous environments. The present study explores specificity in natural populations of a cnidarian-algal model, *Exaiptasia diaphana*, across a latitudinal gradient to understand the genetic and environmental effects driving host-symbiont associations, and their relation to heritable and/or environmental symbiont acquisition. We found that symbiotic associations were extremely flexible in *E. diaphana*, regardless of transmission mode. *E. diaphana* were capable of associating with diverse symbiont communities across genetically identical hosts seeded with vertically transmitted symbionts, as well as across highly connected host populations which acquire symbionts horizontally. Host population connectivity was complex and unrelated to geographic distance, whereas symbiont community composition tracked the thermal gradient, potentially due to context dependent biotic interactions. These results indicate that in a flexible symbiosis, symbiont communities are environmentally-determined, suggesting the future of this symbiosis will likely depend on climate adaptation of symbionts.

## Introduction

Mutualisms are particularly vulnerable to climate change as two or more species must adapt in concert to persist. In many symbioses, generation time and dispersal are de-coupled between hosts and their symbionts [1], posing an evolutionary challenge under a changing environment. Theoretically, specific host-symbiont associations which persist across multiple generations could increase cooperation between partners [2]. However, it may also be advantageous to maintain flexible associations to facilitate acclimatization in the short-term [3]. As the degree of specificity is variable across symbioses [4–6] and will influence the evolutionary trajectory of ecological interactions in future environments [7], it is important to understand how genomic and environmental factors influence host-symbiont associations in contemporary populations.

The diversity of microorganisms in the environment necessitates interaction mechanisms that confer some level of specificity to establish beneficial partnerships and protect against pathogens [8,9]. Host-symbiont associations can be maintained across generations through vertical transmission, where symbionts are directly inherited from parents. Environmental acquisition of symbionts, on the other hand, or horizontal transmission, requires some degree of partner choice for specificity to be maintained. Partner choice during horizontal transmission can be constrained by both genetic and environmental effects including host-symbiont recognition mechanisms, the internal host environment, competition between symbionts, priority effects, and biogeography [10,11]. Horizontal transmitters tend to be more flexible in their associations than vertical transmitters, yet there is evidence of highly specific environmentally acquired symbioses, such as legume-*Rhizobium* and squid-*Vibrio* symbioses, driven by specific host and symbiont traits [5,12]. Though vertical transmission may have evolved as a mechanism to increase specificity between partners, thereby aligning fitness and dispersal [2], vertically transmitting species can also retain the ability to environmentally acquire symbionts, leading to intermediate or mixed transmission modes [13]. Therefore, transmission mode does not necessarily determine specificity, but rather can act as a mechanism to increase specificity by extending the timeframe for coevolution to occur. Understanding the drivers and constraints on symbiont community composition in species with mixed transmission can provide insight into the relationship between symbiont transmission, specificity, and co-evolution.

Here, we explore the genetic and environmental mechanisms driving host-symbiont associations and their level of specificity in an anemone symbiosis with mixed transmission. Commonly known as Aiptasia, *Exaiptasia diaphana* is an emerging model for coral-algal symbiosis [14–17], an ecologically important mutualism that is notoriously threatened by environmental change. Three classes of cnidarians, including coral and sea anemones, host endosymbiotic dinoflagellates in the family Symbiodiniaceae [18], but specificity varies greatly across species (reviewed in [4]), and the mechanisms driving this variation remain unknown. Although transmission mode is related to specificity in cnidarians, there are exceptions which challenge our understanding of how specificity evolves [13,19].

By leveraging a model symbiosis where vertical and horizontal transmission occur during discrete reproductive cycles, we can better understand the mechanisms driving specificity, which is paramount to predicting future stability of thermally sensitive cnidarian-algal symbioses. Aiptasia reproduces both clonally through pedal laceration and sexually by broadcast spawning. During clonal reproduction, symbionts are passed vertically to clonal offspring, which form through tissue budding near the pedal disk [20]. Sexually produced gametes, on the other hand, are aposymbiotic and must acquire symbionts horizontally [21,22]. In the laboratory, horizontal transmission can also be induced in adults, by chemically eliminating symbiont populations and introducing novel symbiont types [23]. However, infection success and maintenance varies across host-symbiont combinations in the lab [21,24–26], implying differences in the degree of host and/or symbiont specificity. In this study, we use 2bRAD and ITS2 amplicon sequencing to profile genetic diversity of naturally occurring Aiptasia and their symbionts across a latitudinal gradient in Southern Florida. By capturing genetically unique hosts over a latitudinal thermal gradient, as well as clonally produced anemones, we were able to explore genetic and environmental effects on symbiont community composition, the capacity for partner specificity, and their relation to transmission mode.

## Methods

### Sample collection

Aiptasia were sampled from six natural populations in Florida spanning a latitudinal gradient from July-September 2019 (see Table S1 for GPS coordinates) under Florida Fish and Wildlife Conservation Commission Marine Special Activity License #SAL-19-2145-SR. Five sites—Otter Key (OK), Virginia Key (MI), Long Key (LK), Tavernier Key (TK), and Big Pine (BP)—were mangrove forests where Aiptasia colonized outer roots within 1 m depth. The sixth site, Stock Island (SI), was a marina where Aiptasia occurred on floating docks. Each site included three subsites ≥30 m apart. At each subsite, five Aiptasia were collected from four roots (≥3 m apart), targeting 60 individuals per population (see Figure 1C for final sample sizes). At SI, five individuals were sampled within ~1 m^2^ areas spaced ≥3 m apart to approximate root-based sampling despite continuity of floating docks. Aiptasia were sampled using a microspatula, transferred to 5 mL tubes and transported on ice to Mote Marine Laboratory (Summerland Key) where they were preserved in 100% ethanol and stored at −20°C. An individual representing the lab strain CC7 was also included as a species control.

**Figure 1.**
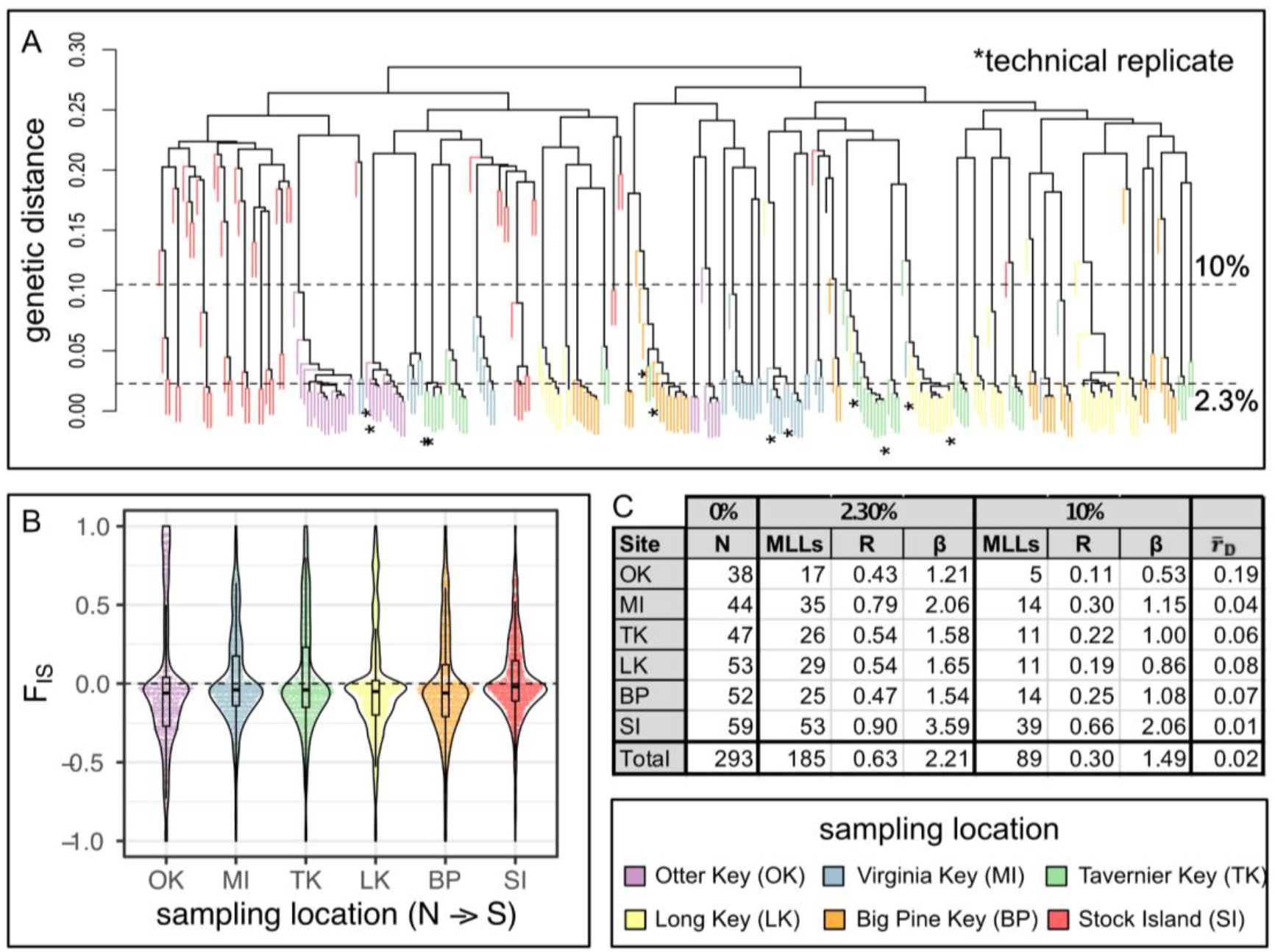
Genet and clonal group identification. A) Hierarchical clustering of pairwise Prevosti distance between samples. Branches represent individual samples colored by sampling location. Asterisks below branches indicate technical sequencing replicate pairs. Two genetic distance thresholds indicated by dashed lines were used for unique genet (>10% distance) and clonal group (<2.3% distance) identification. B) Distribution of per locus inbreeding coefficients (F_IS_) based on sampling location for 1,254 loci. C) Table indicating number of total individuals (N), multi-locus lineages (MLLs) genotypic richness (R), and Pareto β (β) for 2.3% and 10% distance thresholds, and the index of association between loci 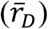.

### Host genotyping

Holobiont genomic DNA was extracted using a Qiagen DNeasy blood & tissue kit. For host genotyping, reduced representation 2bRAD libraries [27] were prepared using a hybrid protocol targeting 1/8^th^ of BcgI sites [28] (https://github.com/z0on/2bRAD_denovo). Library preparation was duplicated for six samples, generating technical replicate pairs for sequencing error estimation and clonal group identification. Libraries were first sequenced on the NextSeq 550 by the USC Genome Core in 2021, yielding ~163k reads per sample on average (51M total). High clonality of this library resulted in low diversity, which reduced read output. To increase diversity and read depth, libraries were pooled with other species and two additional sequencing runs were completed in May 2022 on the NextSeq 2000 by the USC NCCC Molecular Genomics core. Combined these runs yielded a total of 161.7M reads (~510k reads per sample on average). Raw reads are publicly available on NCBI PRJNA1263544.

Bioinformatics were performed on the USC CARC HPC system following the pipeline described in https://github.com/z0on/2bRAD_denovo. First, reads were de-multiplexed based on internal ligation adaptor barcodes, adaptors were trimmed, and PCR duplicates removed using a custom perl script. Sequencing runs were processed separately to remove run-dependent PCR duplicates. Reads were then quality filtered to retain only 99% accurate base calls over 100% of the read using the fastx-toolkit and concatenated across runs resulting in 226.5k de-multiplexed, high-quality reads per sample on average. High quality reads were competitively mapped to a combined *E. diaphana* host genome (Formerly *Aiptasia pallida*, Baumgarten et al., 2015) and four symbiont reference genomes (*Symbiodinium microadriaticum* (ITS2 type A1) *[29], Breviolum minutum* (ITS2 type B1) [30], *Cladocopium goreaui* (ITS2 type C1) [31], and *Durusdinium trenchii* [32] using Bowtie2 [33]. Reads mapping to the host genome were retained for subsequent analysis. Remaining host read depth, coverage, and SNP quality was calculated in ANGSD 0.933 [34] to determine quality filter thresholds. Twelve samples with <20% of sites exhibiting at least 5x coverage were excluded from further analysis, resulting in 294 unique samples and 6 technical replicates remaining. Genotype likelihoods were estimated in ANGSD after filtering to retain high confidence SNPs (-removebads 1 -uniqueOnly 1 -minMapQ 20 - minQ 25 -snp_pval 1e-5) present in ≥80% of samples. Additional filters were also used to exclude SNPs with minor allele frequencies less than 0.05, extremely high depth of coverage (10x number of samples), or strongly deviating from Hardy-Weinberg equilibrium (HWE) (p<1e^−5^).

### Reproductive mode and diversity statistics

For calculating statistics to describe the reproductive mode, ANGSD was rerun on the full dataset with the same filtering and run parameters but this time retaining loci out of HWE with at least 10x coverage across 80% of individuals, resulting in 1,248 sites across 294 samples. The R package hierfstat [35] was used to calculate observed and expected heterozygosity and the inbreeding coefficient (F_is_). As interlocus variation in F_is_ can be an indicator of the reproductive mode [36,37], a Levene Test was used to test for unequal variance across sampling locations. Pairwise comparisons of residual variance were then evaluated using a Tukey post-hoc test. poppr [38], an R package specialized for analyzing partially clonal taxa, was used to calculate the index of association between loci 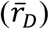 using hard-called genotypes for each site.

### Genet and clonal group identification

Although clones are putatively genetically identical, genotypes could differ due to true biological variation, such as somatic mutations, and/or technical issues, such as genotyping errors and missing data. To account for this, individuals can be collapsed into clonal groups (or multilocus lineages) using a genetic distance cutoff. Pairwise Prevosti distance, which handles missing data, was calculated from genotype calls using the R package poppr [38]. Distance thresholds for collapsing clonal groups were informed by the predicted threshold in poppr using farthest neighbor clustering, as well as genetic distance between technical sequencing replicates. Farthest neighbor clustering was the most conservative, assigning individuals to clonal groups if differentiation was below 2.3%, whereas maximum differentiation between technical replicates was 6.2% (min: 1.6%, median: 3.2%, mean: 3.4%) (Figure S1). We also defined a third threshold (10%) to confidently subsample genetically distinct individuals (genets) and exclude potential clones. This threshold of 10% was chosen based on the frequency distribution of pairwise genetic distance, where the valley between the first and second peak (Figure S2) indicates a threshold above which individuals are considered unique genets [39]. Genotypic richness (R) was calculated by dividing the number of clonal groups (MLLs) by the total number of individuals sampled (R=G-1/N-1 [40]) across 2.3% and 10% distance thresholds. The Pareto index (β), calculated as the negative slope of the Pareto distribution of clonal group size [41], was also calculated across distance thresholds using the “poweRlaw” package in R [42].

### Host population structure

Population structure was evaluated for the complete dataset without filtering individuals and pruned datasets after removing potential clones (greater than 2.3% or 10% differentiation). For all thresholds, loci strongly deviating from HWE (p<1e^−5^) were removed and ngsLD [43] was used to filter physically linked loci (r2 > 0.1 within 50kb distance). Admixture was evaluated from genotype likelihoods using NGSadmix [44] for 1-7 genetic clusters (k) and 20 independent runs were generated for each k. Delta k was calculated following Evanno et al. (2005) to determine the number of source populations (k). As all datasets supported an optimum K of 2 (Figure S3-S8), we opted to present the admixture results and subsequent analyses using the greatest distance threshold (10%), in order to remove putative clones and match the mathematical assumptions of the population genetic models. After filtering, 89 unique genets and 478 loci remained.

Genetic differentiation was evaluated by calculating pairwise F_st_ values between populations from hard-called genotypes using hierfstat [45]. Scaled F_st_ values were plotted against geographical distance (mean over water distance measured in Google Earth®) and to evaluate isolation by distance (IBD) using the function mantel.rtest from the ‘ade4’ package [46] in R with 999 permutations. Linear models were also used to analyse location specific patterns of genetic and geographic differentiation. Migration rates between populations were modeled for multiple deme sizes (200, 300, 400) with 5M MCMC iterations and a 2M burn-in period using EEMS (estimated effective migration surfaces) [47]. Model convergence and migration rates were visualized using the associated R package rEEMSplots [47].

### Symbiont community profiling

Symbiont DNA was simultaneously extracted with host DNA as described above. Amplicon libraries were prepared by amplifying the ribosomal ITS2 region (x20-30 cycles) using the SYM_VAR_5.8S2/SYM_VAR_REV primers [48–50], followed by a second PCR (x6 cycles) to add sample-specific barcodes. Paired-end 250bp reads were sequenced on the MiSeq v2 by the USC NCCC Molecular Genomics core with 30% PhiX spike-in which yielded a depth of 64.1M reads across two runs (nano and v2 chemistry) in June and July 2022. Forward and reverse reads were concatenated across runs (~98.3k paired reads per sample on average) and analyzed in SymPortal using the default settings. To verify profiles for samples with high cycle numbers (>25), 39 sample libraries were re-prepared from remaining DNA with the inclusion of negative controls and re-sequenced on the MiSeq v2 by the USC NCCC Molecular Genomics Core in December 2022. Re-sequenced samples and the negative control were analyzed in SymPortal, and confirmed that the original symbiont profiles represent true biological signals (Figure S9).

ITS2 profiles were generated in SymPortal by collapsing co-occurring defining intragenomic variants (DIVs) present in at least four individuals (Hume et al., 2019). To explore whether clonality increased the number of unique profiles, SymPortal was rerun including only unique genets (Figure S10). Symbiont profile absolute abundance was imported into the R package Phyloseq [51] to generate relative abundance plots. Flexibility in symbiont community composition among clones was evaluated at symbiont profile, type, and genus levels by exploring variation in dominant community members (>50% relative abundance) across clonemates using the most conservative threshold (host genetic distance <2.3%). Community composition and beta diversity was compared across subsites and sites through PERMANOVA and PERMDISP tests in the R package vegan [52]. Differences in community composition (bray-curtis dissimilarity) were correlated to spatial distance, temperature differences, and host genetic differentiation (F_st_) across sites using mantel tests from the vegan package to determine the factors driving symbiont community composition at genus, ITS2 type, and profile levels. To evaluate co-phylogeny between hosts and symbionts, unifrac distance between symbiont profiles was correlated to Prevosti distance between host genets using the Procrustean Approach to Cophylogeny (PACo) package in R which normalizes the scale across phylogenies [53]. A binary matrix was generated to represent the presence or absence of each symbiont profile detected within unique host genets (genetic distance >10%) to indicate host-symbiont associations. The phylogenetically-informed interaction network was statistically compared to randomly generated interactions across 1000 permutations. Host-symbiont associations were tested against multiple randomisation algorithms to detect co-divergence where the symbiont tracks host phylogeny (r0), the host tracks symbiont phylogeny (c0), or it is unclear which group is tracking the other (swap and backtrack) [53].

## Results

### Host reproductive mode

Genotypic diversity (R) and the distribution of clonal membership (Pareto β) were consistently lowest for Otter Key (OK) and highest for Stock Island (SI) across all genetic distance thresholds (Figure 1C). Stock Island (SI) also had the lowest index of association (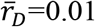, Figure 1C) and significantly less variation in per locus F_is_ estimates compared to all other sampling locations (Table S2;Figure 1C). Pareto β was greater than 2 regardless of genetic distance threshold (3.59 > β > 2.06, Figure 1C), suggesting the SI population is primarily sexual [54]. In contrast, Otter key (OK) had the lowest genotypic diversity (R=0.43) and distribution of clonal membership (1.21 > Pareto β > 0.53), indicating this population likely has a much higher rate of clonality [54]. This is further supported by OK having the greatest index of association 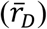 and significantly more variation in per locus F_is_ estimates compared to all other sampling locations (Table S2; Figure 1BC). Other populations (MI, TK, LK, and BP) had intermediate values of Pareto β (Figure 1C), suggesting partial clonality (0.7 < β < 2, [54]). Together these results indicate that natural populations of Aiptasia are partially clonal, and the extent of clonal reproduction varies among locations.

Under the threshold for conservatively assigning clonal groups (genetic distance <2.3%), clonemates were found on up to four roots within a site (Table S3), and occasionally across subsites (Table S4), indicating local migration of adults or clonal offspring. However, genotypic diversity was also present across small spatial scales. When considering only high confidence, unique genets (genetic distance >10%), mangrove roots were colonized by 1-2 genets on average, with up to four genets colonizing a single root (Figure S11, Table S5). At the marina (SI), seven unique genets were identified in a ~1m^2^ area on average (Figure S11, Table S5), highlighting the greater genotypic diversity of this site. Notably, clonal migration and genotypic diversity increase with more lenient thresholds (lower or higher genetic distance respectively) so reported values likely represent the lower limits of these processes.

### Host population structure

Based on the admixture analysis, two genetic clusters (K=2) were detected across all clonality thresholds (Figure 2A; Figure S3-S8). The most prominent genetic break was between SI and all other populations (Figure 2A), which was consistent with an MDS analysis of IBS distance (Figure S12). At least one anemone from each site (23 out of 51 genets not sourced from SI) shared greater than 25% ancestry with those from the SI population, suggesting admixture between subpopulations. Despite subpopulation structure, gene flow was high overall between populations (0.01 > F_st_ < 0.07, Figure 2C, S13). F_st_ was lowest (0.01) between sites less than 100 km apart, with the exception of BP and SI which had one of the highest F_st_ values (55.12 km, F_st_ = 0.06). In support of this, there was high effective migration in the upper and middle keys, but a potential genetic break in the lower keys (Figure 2B). As estimating effective migration surfaces can only indicate possible barriers to gene flow and not directionality, we additionally performed an isolation by distance analysis to test whether geographic distance linearly scaled with genetic differentiation. Although no isolation by distance (IBD) was detected across all locations, we found a negative correlation between genetic and geographic distance when comparing SI and OK to other sites, opposite to what is expected under IBD. Specifically, there was a significant negative correlation between geographic and genetic distance of SI and all other Florida Keys sites (p=0.02, R^2^=0.9375, Figure 2C – orange line). Additionally, MI had the least amount of differentiation compared to OK (F_st_=0.04), despite being the most geographically distant sites (Figure 2C, S13), and there was also a significant negative correlation between OK and all other sites (p=0.001,R^2^=0.9712, Figure 2C – gray line). No significant positive or negative associations between genetic and geographic distance were found among upper and middle keys sites.

**Figure 2.**
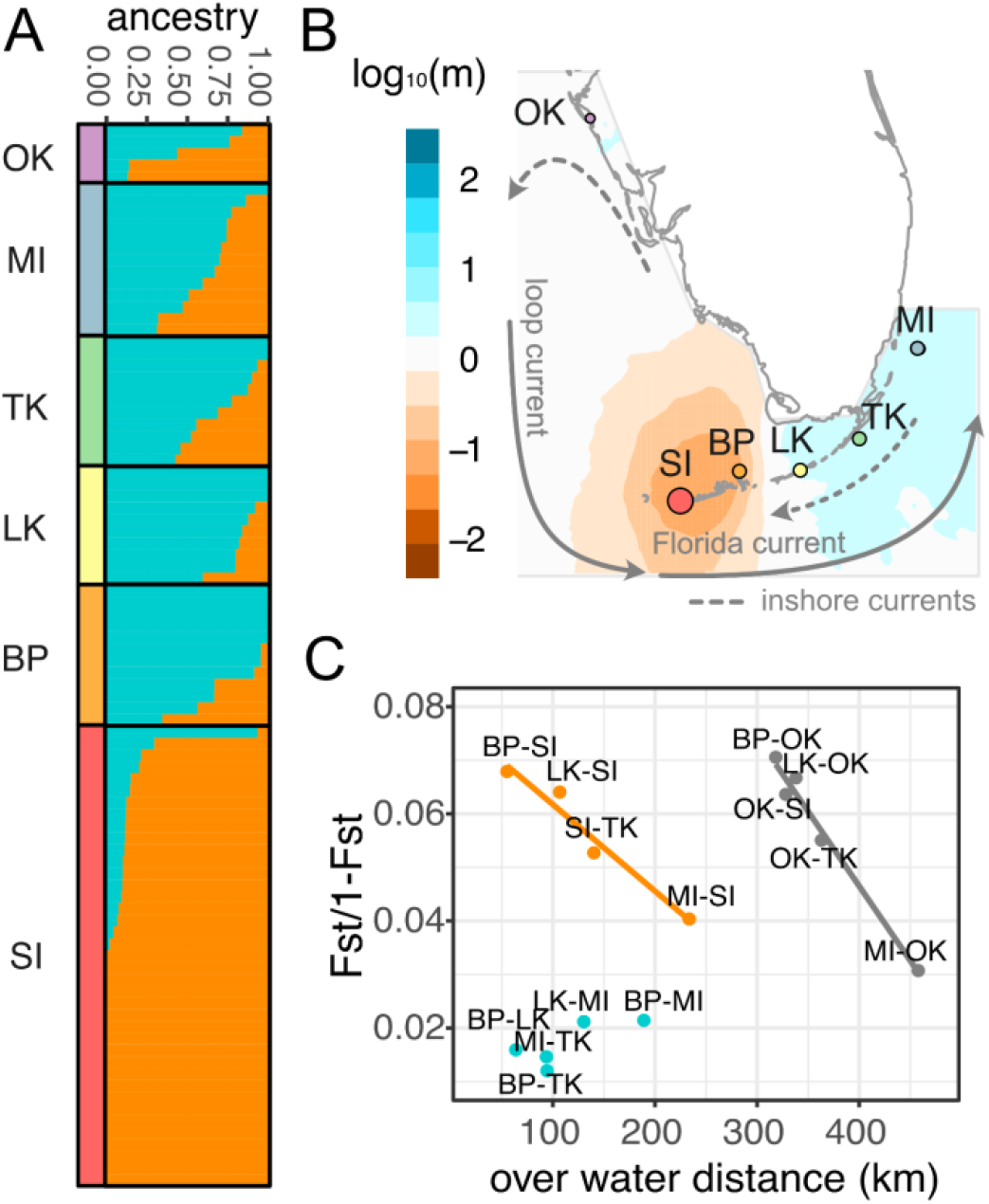
Host genetic differentiation based on geographic distance. A) Admixture proportions of each individual (bars) grouped by sampling location. B) Estimated effective migration rates between sites. Colors indicate greater (blue) or less (orange) migration than would be expected under IBD. Arrows indicate predominant directionality of sea surface currents following [55,56]. C) Pairwise geographic distance and genetic differentiation between sites. The orange and gray lines indicate a significant negative relationship between geographic distance and genetic differentiation for pairwise differences across SI and OK, respectively. Points labels represent pairwise comparisons between Otter Key (OK), Miami (MI), Tavernier Key (TK), Long Key (LK), Big Pine Key (BP), and Stock Island (SI).

### Symbiont community composition and distribution

Thirty-five symbiont profiles comprising representatives of four *Symbiodiniaceae* genera (*Symbiodinium, Breviolum, Cladocopium*, and *Durusdinium*) were identified as symbiotic partners of naturally sourced Aiptasia (Figure 3). *Symbiodinium* was the most abundant and diverse genus, with 77% of samples being dominated by one of 13 *Symbiodinium* profiles. *Breviolum* and *Cladocopium* were less abundant, but relatively diverse, with 11 and 9 unique profiles, respectively. *Durusdinium* was the least abundant genus and was only detected in two samples at background levels (<10% relative abundance). The majority of individuals (78%) hosted a single symbiont profile whereas the minority (22%) hosted mixed symbiont communities composed of more than one symbiont genus.

**Figure 3.**
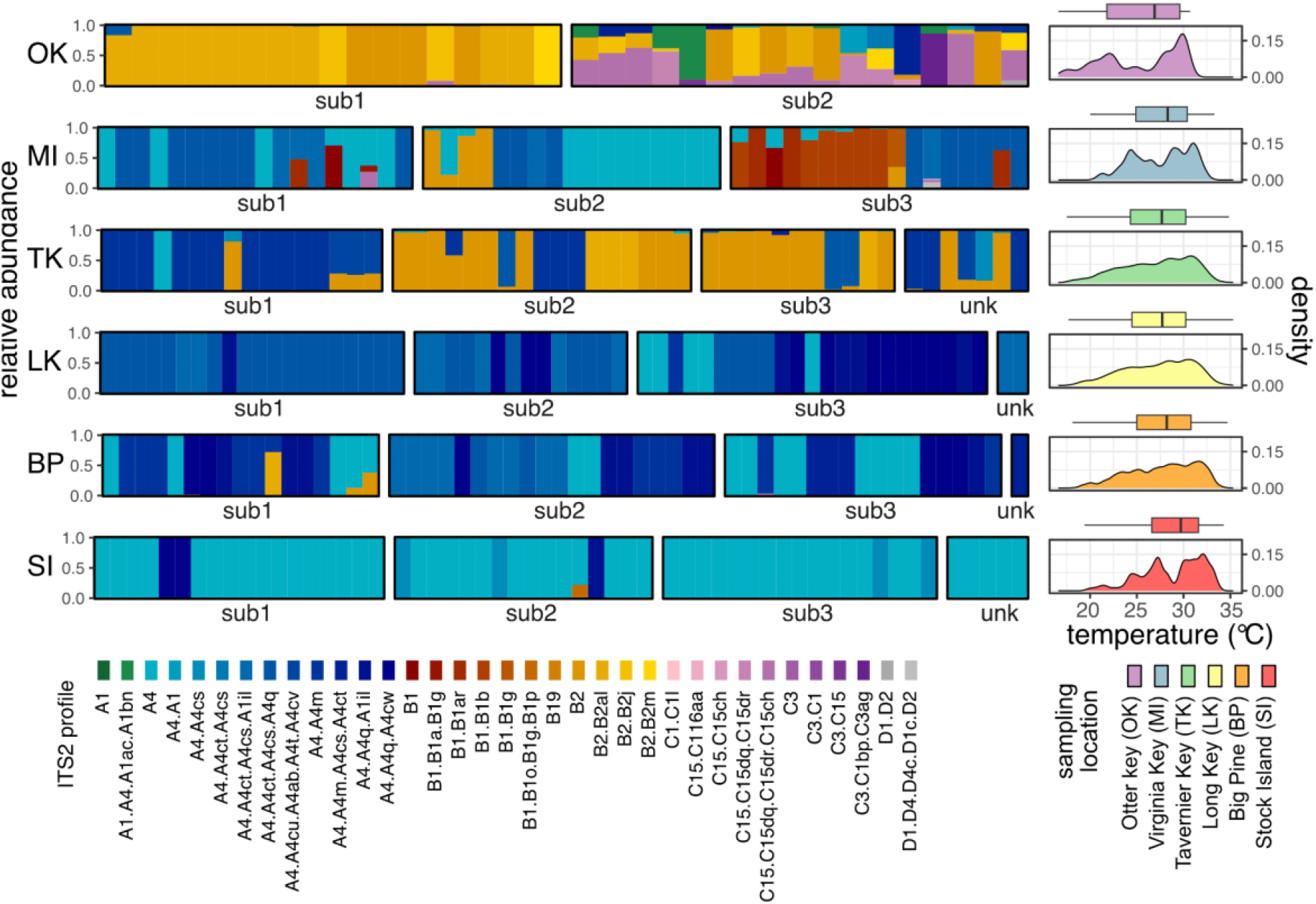
Relative abundance of symbiont profiles (left) and temperature profiles (right) across sites. Each bar represents an individual sample. Samples are grouped by subsite (bolded boxes) within sampling location (rows) ordered from north (top) to south (bottom). “Unk” indicates a sample from an unknown subsite. Symbiont color families represent genus-level designations and gradation within color families denote unique ITS2 profiles. Density plots of temperatures from nearby buoy data (right) are grouped and colored by sampling location.

Symbiont genera and ITS2 types significantly differed between northern and southern sites (PERMANOVA, p=0.001; Table S6-7). *Cladocopium* and *Breviolum* symbionts more frequently dominated Aiptasia in northern sites but were uncommon or absent in lower latitude populations (Figure 3). *Cladocopium* symbionts were restricted to the two northernmost sites, and only dominant in Otter Key individuals. Otter Key Aiptasia were also frequently dominated by *Breviolum* profiles, but rarely by *Symbiodinium* profiles (2 individuals). In the upper keys (MI and TK), both *Breviolum* and *Symbiodinium* frequently dominated Aiptasia. However, *Breviolum* was uncommon south of Tavernier Key (Figure 3), and southern individuals (LK, BP, and SI) were exclusively dominated by *Symbiodinium* A4 profiles, resulting in greater genus and ITS2-type diversity in northern (OK, MI, TK) versus southern sites (Table S9-10). However, despite little variation in symbionts at the genus and ITS2 type-level within southern sites, *Symbiodinium* profiles significantly differed (pairwise adonis, p=0.001; Table S8) and beta diversity was significantly lower in the SI population in comparison to other southern sites (permutest, p<0.001, Figure S14, Table S11).

Differences in temperature across sites was significantly correlated with differences in symbiont community composition at the genus and ITS2-type levels (mantel test, genus: r=0.6467 p=0.029167, ITS2 type: r=0.6725 p=0.027778). Though spatial distance between sites was not significantly correlated with differences in symbiont community composition (mantel test, genus:, r=0.4957 p=0.16389, ITS2 type: r=0.5284 p=0.16389, ITS2 profile: r=-0.1381 p=0.56806), the relative abundance of individual ITS2 profiles exhibited significant spatial autocorrelation (Moran’s I: min=0.09, max=0.60, mean=0.38, Table S12). After removing putative clones to account for vertical transmission, only six ITS2 profiles had high enough frequency to test for spatial autocorrelation. Three of these six profiles showed significant spatial autocorrelation, though correlation coefficients were slightly weaker compared to the dataset including clones (Table S13).

### Host-symbiont specificity and co-divergence

Symbiont fidelity was observed in 22 clonal groups (48%) wherein all clones hosted an identical symbiont profile. Conversely, symbiont flexibility was observed in the other 24 (52%) clonal groups, where dominant symbiont profiles varied among clonemates (Figure 4B, Figure S15). Of these 24 host clonal groups with flexible associations, 13 exhibited symbiont differences at the profile-level, 1 at the ITS2 type-level, and 10 at the genus-level (Figure 4B, Figure S15, clonality threshold <2.3%). Host clonal groups exhibiting symbiont flexibility were observed across subsites and even along individual mangrove roots.

**Figure 4.**
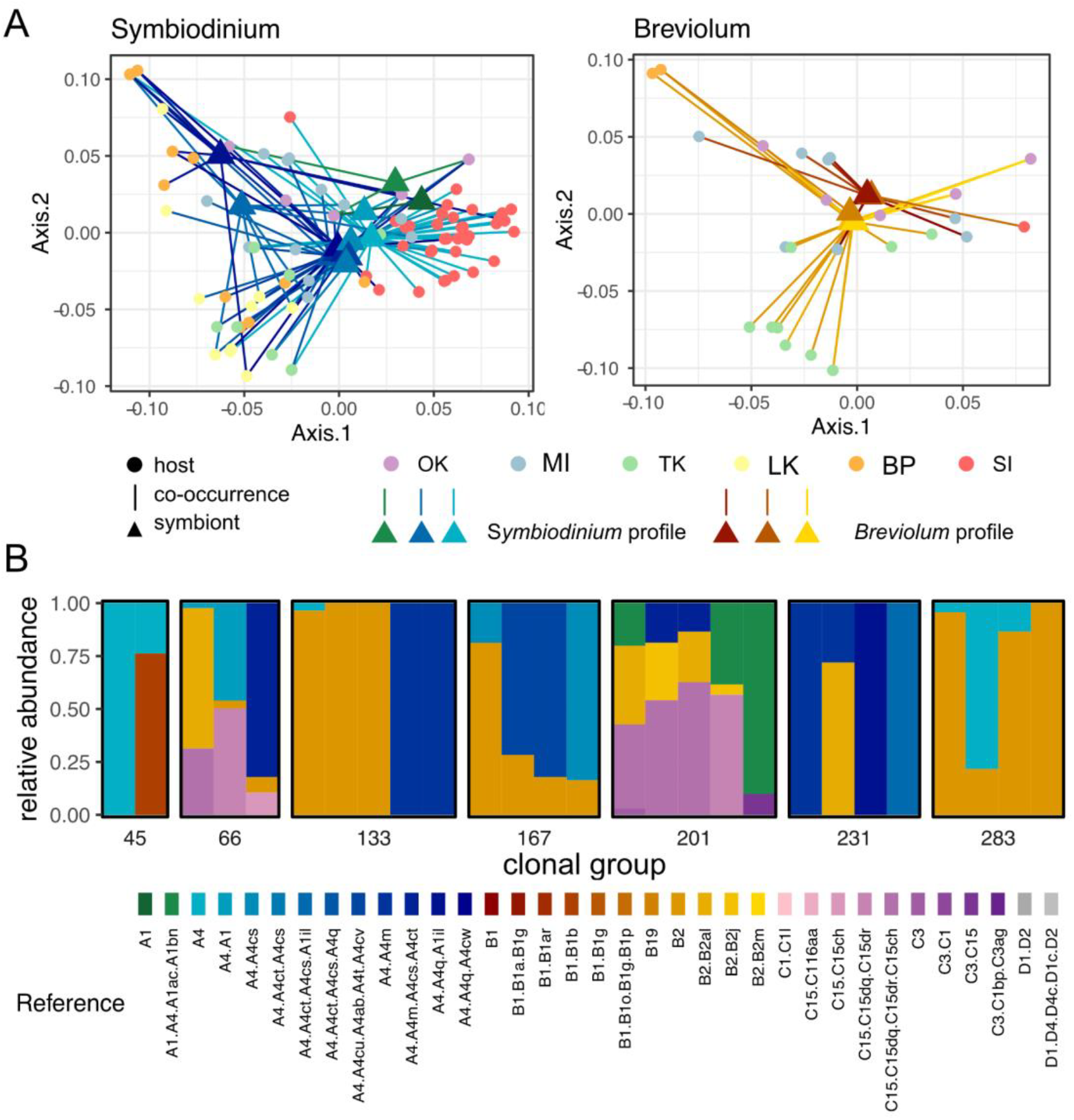
Co-divergence and host flexibility. A) Symbiont to host co-occurrence networks (lines) based on host (circles colored by sampling location) and symbiont (triangles colored by major ITS2 type) genetic divergence within genera (left:Symbiodinium, right:Breviolum). B) The relative abundance of symbiont profiles within host clonal groups which exhibit flexibility in dominant symbiont genera across individuals. Each column represents a single individual and clonemates are binned within bolded boxes.

After excluding clones to consider only unique genets with horizontally acquired symbionts (genetic threshold >10%), host genetic differentiation between sampling locations (Fst) was not significantly correlated with differences in symbiont community composition at the genus, ITS2 type, or profile level (Mantel test, genus: r= 0.4221 p=0.065278, ITS2 type: r=0.4112 p=0.0875, ITS2 profile: r=0.4137, p=0.14444), suggesting horizontally acquired symbiont communities at the population level are not driven by host genetics. However, when considering pairwise genetic distance between individual host genets and ITS2 profiles, host genetic distance was weakly correlated with symbiont profiles in some genera (Mantel test, *Symbiodinium* r=0.15, p=0.001, *Breviolum* r=0.173, p=0.001, *Cladocopium* r=0.1563, p=0.071), and even after removing clones in *Symbiodinium* (Mantel test *Symbiodinium* r=0.175, p=0.005, *Breviolum* r=0.1583, p=0.108, *Cladocopium*: r=0.8361, p=0.25). This is further supported by significant co-divergence between host and symbiont co-occurrence networks within symbiont genera (PACo r0, *Symbiodinium*: m2xy=5.32, p=0; *Breviolum*: m2xy=1.79, p=0.002, Figure 4), indicating the potential for host-symbiont associations is partly aligned with host and symbiont genetics. We considered null models which randomized symbiont connections (c0), host connections (r0), or both (swap and backtrack) [53]. Co-divergence of *Symbiodinium* and *Breviolum* networks were significantly different than null models where either host or symbiont could track the other (swap and backtrack) and where symbiont tracked host divergence (r0), but not for the model where the host tracked symbiont divergence (c0) (Table S14). *Cladocopium* divergence was not correlated to host genetic distance (Table S14), though profiles in this genus were not frequently present in Aiptasia communities.

## Discussion

Though specificity of host-symbiont associations and coevolution could optimize partner benefit, flexibility may be especially important for acclimatization when partners are likely to encounter variable environments within and across generations. In this study, we found that both Aiptasia and its symbiotic algae were capable of association with genetically diverse partners, highlighting the flexibility of this symbiosis. Although hosts and symbionts did exhibit signatures of co-divergence, this pattern was based on occurrence and did not consider community structure within individuals. Symbiont community composition in-hopsite was locally determined and tracked the thermal gradient, suggesting that symbiont communities are driven by the environment, rather than host genetics. In support of this, Aiptasia were capable of associating with divergent partners, even within clonal groups seeded with vertically transmitted symbionts. At a population level, hosts exhibited high gene flow among sampling locations and differentiation did not track the thermal gradient, which would likely constrain thermal adaptation of the host. Therefore, associating with locally adapted symbionts may allow Aiptasia to acclimatize to different environments, favoring flexibility in the symbiotic association.

Together these results indicate that in this flexible symbiosis, symbiont communities are environmentally-determined, despite alignment of dispersal during vertical and possibly horizontal transmission, suggesting the future of this symbiosis will likely depend on climate adaptation of symbionts.

### In-hopsite symbiont communities are driven by local environments

The latitudinal succession of symbiont genera observed in the present study (Figure 3) aligns with prior work on the ecological and physiological differences among Symbiodiniaceae genera. For example, *Cladocopium* are consistently found in deeper and more turbid environments [57–59] and at higher latitudes [60–64]. *Symbiodinium* also has greater thermal tolerance than *Breviolum* in culture [65], which could explain the restricted distribution of *Cladocopium* and *Breviolum* to cooler, northern sites observed here. Although symbiont communities could be reflective of partner availability, a population of Aiptasia in the Florida Keys was recently shown to exclusively host *Symbiodinium* despite diverse environmental symbiont pools including *Cladocopium* and *Breviolum* species [66]. In support of this, experimental manipulations of Aiptasia in the laboratory have demonstrated that *Breviolum* preferentially infected Aiptasia over *Symbiodinium* under ambient conditions (25C), but infection dynamics were reversed at elevated temperature (32C) [26]. Together this suggests that competition during host colonization likely explains thermal structuring of symbiont communities rather than environmental filtering. However, symbiont species and strains within genera can also have diverse physiology [65], so environmental sampling of symbiont pools coupled with physiological assays are needed to resolve the extent to which community structure is influenced by the external environment and/or its interaction with the host environment.

Divergent symbiont communities among clones with vertically transmitted symbionts also indicates an influence of microhabitat on symbiont community composition *in hospite*. Clonal groups with divergent symbiont genera were detected across subsites and roots, and even along the same mangrove root (Figure 4C). Although we do not have micro-scale environmental data, individual corals can have spatially partitioned symbiont community members based on colony microhabitats [57,67], supporting that subtle changes in the environment can affect symbiont community structure. Additionally, shuffling or switching dominant symbiont community members within individuals has been identified as an acclimatory mechanism, facilitating symbiont associations adapted to novel environments. Although many observations follow disturbance events such as coral bleaching, symbiont switching/shuffling can also occur seasonally in the absence of dysbiosis [68,69]. Further, Aiptasia symbiont communities undergo population bottlenecks during pedal laceration followed by rapid proliferation as hosts develop, which could give low abundance or novel symbionts a chance to outcompete poorly adapted symbionts.

Our results indicate that Aiptasia symbioses may be resistant to climate change based on their ability to switch/shuffle symbionts across different environments to locally adapted symbionts. In corals, shuffling of communities to heat tolerant members can increase coral thermal tolerance by 1-1.5°C [70]. Additionally in models of coral evolution, symbiont switching/shuffling was more effective in increasing coral survival compared to symbiont adaptation alone, but both were required for persistence under the most extreme climate projections [71]. Consistently, experimental evolution is a proposed intervention strategy to generate more thermally tolerant symbionts for inoculatating coral [72]. However, most preliminary studies have focused on environmental acquisition of heat-evolved symbionts during early life stages (larvae and juveniles), and the capacity for adults to switch or shuffle to evolved symbionts in the absence of an induced bleaching disturbance is unknown. As Aiptasia symbiont communities can be easily manipulated in the lab along with the external environment during both juvenile and adult stages, Aiptasia represents an important model system to study both natural and experimentally-evolved symbiont communities within and across generations and environments.

### Host and symbiont dispersal is aligned during horizontal transmission

Evolutionary theory predicts that horizontally transmitted symbionts would show limited co-phylogeny, as selection pressure would not be consistent for symbiotic and free-living stages. This theory is consistent with empirical evidence of other horizontally transmitted symbioses, including many animal-bacterial symbioses [10,73,74]. However, several studies of coral symbionts have identified patterns of co-divergence in spite of horizontal transmission [75,76], consistent with patterns observed here. Horizontally transmitted Vibrio symbionts also exhibit co-divergence with squid hosts [77], which is in part driven by both specificity and enrichment of local environments with parental symbionts [78]. Given the flexibility of both host and symbiont partners observed in this study, patterns of co-divergence in Aiptasia are most likely driven by similar dispersal of partners leading to co-occurrence.

Both hosts and symbionts exhibited broad dispersal across the studied range. Host population structure indicated low genetic differentiation overall that was not explained by geographic distance, suggesting high dispersal capacity. Broadcast spawning hosts typically have long pelagic larval duration, which facilitates broad dispersal and commonly leads to low genetic differentiation between geographically distant populations [79]. Aiptasia larvae can survive up to 8 weeks in the lab (personal observation), providing ample time for transport between distant locations. Although symbionts are assumed to have more restricted dispersal due to their sedimentary nature and poor swimming capacity [80], existing work on population genetics of Symbiodiniaceae has reported large dispersal ranges [3,81], consistent with patterns observed here.

Marine larvae are generally transported through sea surface currents [82–85], which could explain genetic differentiation of Aiptasia. For example, counterclockwise offshore transport via the Mexico Loop and Florida currents [86,87] could be facilitating high gene flow between OK/SI and MI hosts (Figure 2B, 2C). Similarly, in *Symbiodinium*, co-divergence seems mainly to be driven by co-occurrence of A1 and A4 ITS2 types with Otter Key, Miami, and Stock Island hosts, whereas other A4 variants were more likely to co-occur with hosts originating from the middle keys (LK, TK, BP) (Figure 4A). Given existing evidence of broad dispersal of symbionts by sea surface currents [81], hosts and symbionts may be co-transported, leading to co-divergence. Alternatively, Aiptasia larvae can also uptake symbionts at the larval stage, so could be indirectly dispersing symbionts from source populations. However, it is important to note that environmental symbiont pools were not sampled in this study and genetic distance of symbiont profiles may not accurately represent population differentiation of symbionts. Population genetic studies of symbionts in- and ex-hospite are needed to fully resolve the mechanisms of symbiont dispersal.

### Mixed transmission promotes host flexibility

Despite the broad dispersal capacity of symbionts, the relative abundance of symbiont profiles was driven by thermal environments. Flexibility of the host may therefore be key to establishing symbiosis with locally adapted symbionts, especially when offspring are unlikely to be retained within parental populations. Previous studies on symbiont communities in Aiptasia have reported both specific and flexible host populations [88,89], with populations from Florida exhibiting more flexibility compared to globally distributed populations which exclusively associate with ITS2 type B1 symbionts [89]. Although we also observed differences in symbiont community diversity between host populations (SI vs others), the SI population primarily hosted A4 symbionts. Notably SI anemones were also sampled from a unique substrate (floating docks), which were warmer and less environmentally heterogeneous compared to mangrove habitats, so it is unclear whether this pattern is driven by specificity, restricted dispersal, or the environment. Importantly host strains from global populations can also be dominated by symbionts other than B1 in the laboratory depending on temperature [26], supporting broad symbiont recognition mechanisms in Aiptasia and importance of environmental conditions in determining community composition.

Corals with horizontal transmission generally exhibit greater flexibility in symbiotic partners compared to vertical transmitters, which leads to a more dominant effect of the environment on symbiont community composition [60,75]. However, there is a growing body of evidence that many vertical transmitting coral do not exclusively inherit symbionts and can also acquire them from their environment [13,19]. This can lead to false patterns of specificity whereby vertical transmitters appear to be exclusive under stable environmental conditions, but new associations are possible following environmental disturbance [90]. Although it is still likely that symbiont communities in horizontal transmitters are more influenced by the environment than those with the capacity for vertical transmission, these patterns argue that primary and secondary ecological succession may be more important in driving symbiotic associations, rather than strict specificity, when environmental acquisition is possible.

Observed specificity during vertical transmission could be due to primary ecological succession such as priority effects, where vertically transmitted symbionts would have an advantage over late arriving, horizontally acquired symbionts. This could lead to stability in associations when environments are consistent. Within clonal groups with vertical transmission, we found retention of symbiont communities occurred about 50% of the time, possibly explained by priority effects. However, disturbance events create the opportunity for secondary succession, where ecological outcomes could differ based on new environmental conditions, and could be leading to community turnover in the other 50% of clonal groups (Figure 4C). Therefore, environmental heterogeneity could constrain the evolution of specificity during vertical transmission due to inconsistent fitness landscapes. However, it is also important to note that vertical transmission in Aiptasia only occurs during clonal reproduction. The evolution of specificity could therefore be less efficient in Aiptasia compared to other systems where vertical transmission occurs during sexual reproduction. Nevertheless, this study challenges the assumption that vertical transmission increases specificity, and emphasizes the need for incorporating diverse environments in future investigations

### Limitations and future recommendations

Although the overall patterns presented here were robust across genetic distance thresholds, diversity metrics could not be absolutely determined due to challenges in classifying genotypic and phylogenetic variation. Genotypes within clonal groups are not always identical and could differ due to true biological variation, such as somatic mutations while still belonging to the same clonal lineage, and/or technical issues, such as genotyping error compounded by missing data. Therefore, genotypes are often collapsed into clonal groups or multilocus lineages using a genetic distance threshold based on the distribution of pairwise genetic distance [38,39] or distance between technical sequencing replicates [91], but both are rarely explored. In the present study, distribution-based methods, such as poppr [38], indicated a lower threshold for collapsing clones, which did not collapse simulated clones (technical replicates) into clonal groups (Figure 1). As low coverage, reduced representation genotyping such as 2bRAD used here tend to have greater levels of sequencing error and missingness [27], this method may be too coarse to resolve true clones. Future studies should prioritize methods that increase the recovery of a genotype at each locus, such as microsatellites [92] or HiPlex SNP genotyping [93], to minimize technical issues and maximize analytical power to detect clonal lineage. Moreover, temporal sampling can help resolve variation in the reproductive mode by comparing genotype frequencies (see [94] for methods and [95] as an example).

Similarly, ITS2 amplicon sequencing is widely used for profiling algal symbiont communities, but interpretations of diversity are challenging (reviewed in [96]). In Symbiodiniaceae, ITS2 is a multi-copy gene, making it difficult to resolve intra-genomic from inter-genomic variation. Collapsing ITS2 types into distinct profiles as done in the present study may be a way to resolve this, but it is unclear whether ITS2 profiles represent diversity at the strain or species level [97]. Further, ITS2 sequences do not align well across symbiont genera, so phylogenetic distance is not factored into diversity estimations when multiple genera are present. Despite these challenges, ITS2 is still important as the first step in describing symbiont community composition when diversity is unknown [96], but analysis of microsatellite and single copy genes are necessary to validate population and community-level results presented here.

## Data availability statement

Trimmed and quality filtered 2bRAD sequences are archived on NCBI (PRJNA1263544) and will be made publicly available upon publication. Symbiont ITS2 amplicon sequences will be made publicly available via the SymPortal database (20220805T211857_mruggeri) upon publication. All scripts for analyses and metadata information are archived and publicly available on github (https://github.com/mruggeri55/AipFLkeysPopGen).

## Acknowledgements

We would like to acknowledge Wyatt Million for field assistance during sampling. Funding for this study was provided by the University of Southern California to CDK.

