## Supplemental document 1 for "Environmentally-determined symbiont communities highlight flexibility of Aiptasia-algal symbiosis"

**Supplemental information for Environmentally-determined symbiont communities highlight flexibility of Aiptasia-algal symbiosis**

Maria Ruggeri[1*], Connie S. Machuca[1], Samuel A. Bedgood[2], Stacy A. Krueger-Hadfield[3], Carly D. Kenkel[1]

[1] Biological Sciences, University of Southern California, Los Angeles, CA 90089

[2] Biological Sciences, University of California, Irvine, CA 92697

[3] Virginia Institute of Marine Science Eastern Shore Laboratory, Wachapreague, VA 23480

### Supplemental figures

Figure S1. Distribution of the number of clonal groups (multi-locus lineages) called over various genetic distance thresholds. Dashed horizontal lines represent distance between technical sequencing replicates. The red line indicates suggested cutoff for clonal groups determined from Poppr models.

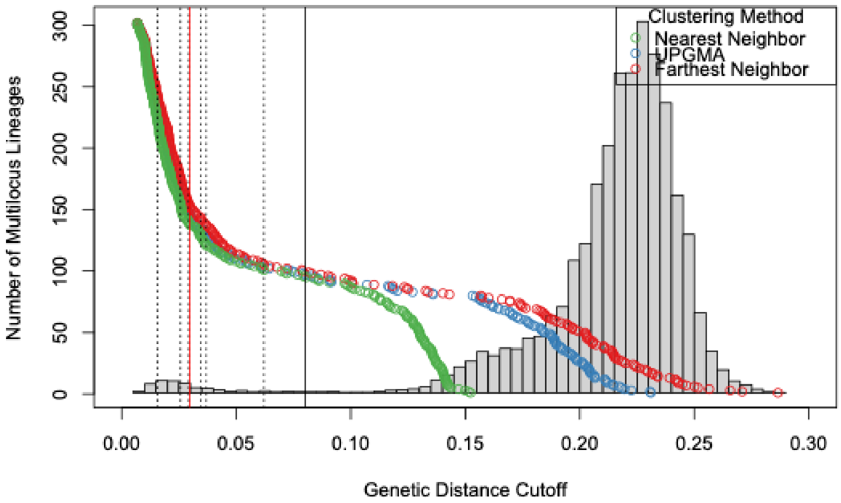


Figure S2. Subset of distribution of the number of clonal groups (multi-locus lineages) called over genetic distance thresholds below 10% indicating a valley between first and second peaks used to define a conservative threshold for population genetic analyses. Dashed horizontal lines represent distance between technical sequencing replicates. The red line indicates suggested cutoff for clonal groups determined from Poppr models.

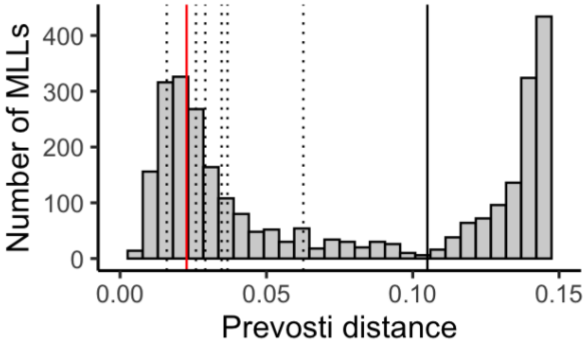

Figure S3. Admixture proportions for various Ks including only high confidence unique genets (genetic distance >10%).

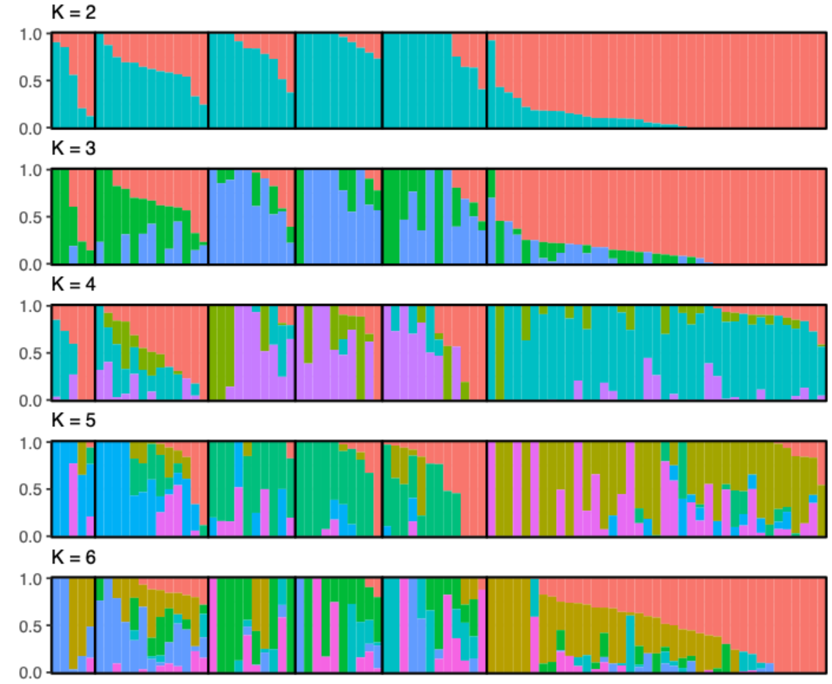


Figure S4. DeltaK MLLs89

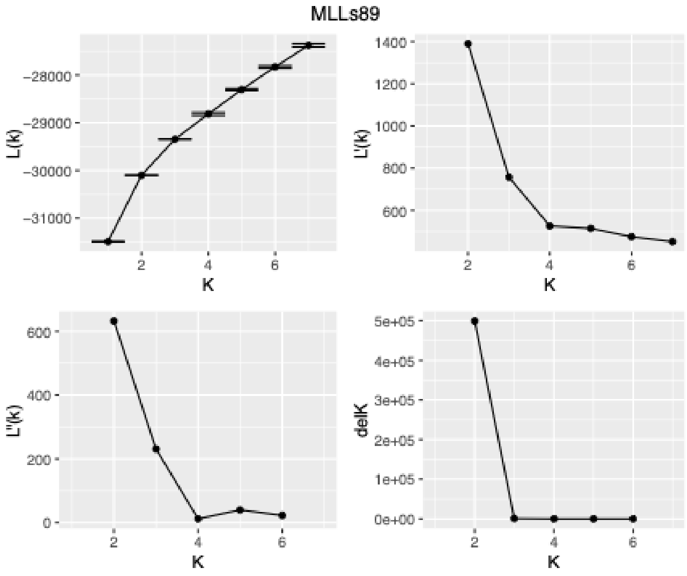

Figure S5. DeltaK MLLs184

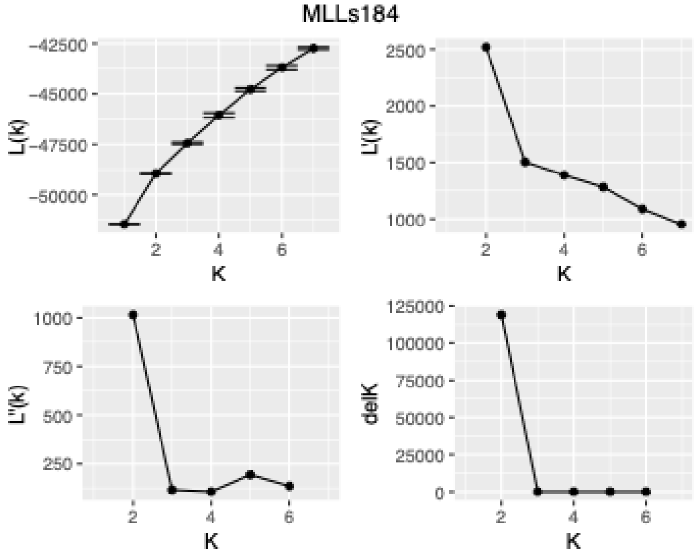


Figure S6. DeltaK all individuals sampled

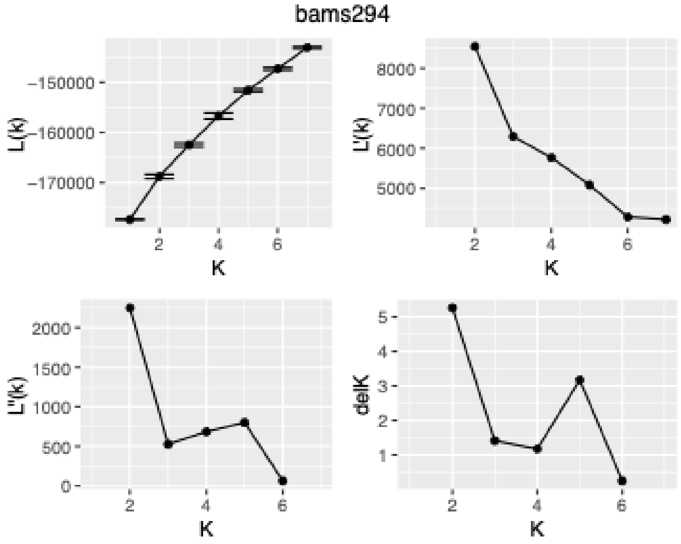


Figure S7. K of 2 admixture proportions for all individuals samples ordered by sampling location.

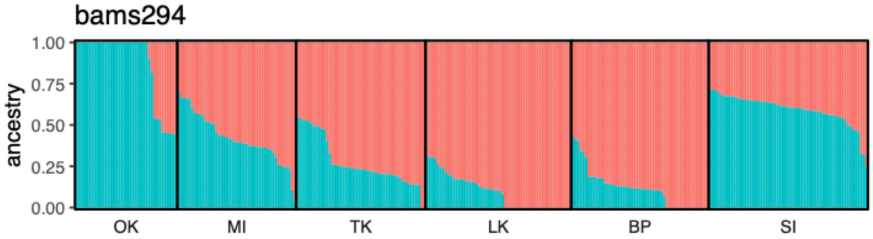

Figure S8. K of 2 admixture proportions for after pruning putative clones (genetic distance <2.3%) order by sampling location.

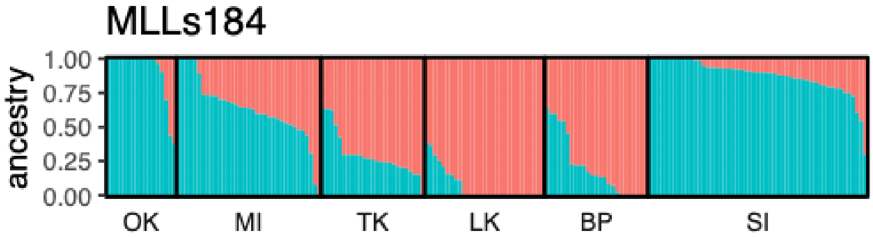


Figure S9. Absolute abundance of re-sequenced samples including three negative controls (NC1-3). Colors indicate symbiont genera (*Symbiodinium* – pink, *Breviolum* – green, *Cladocopium* – blue, and *Durusdinium* – purple).

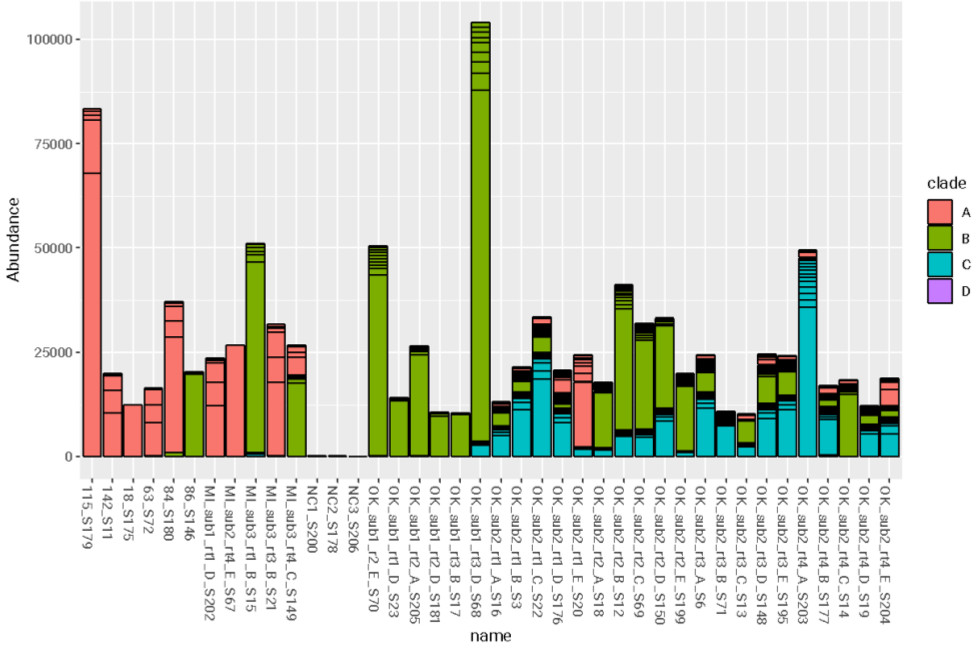

Figure S10.Relative abundance of symbiont profiles analyzed with (top) and without (bottom) clones.

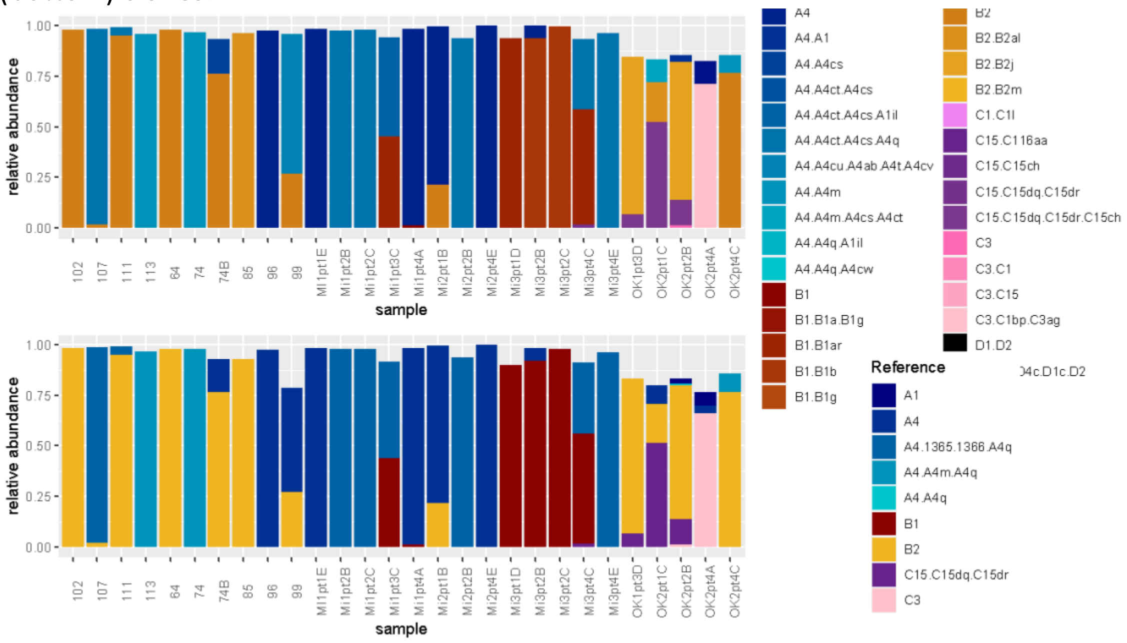

Figure S11.IBS dendrogram within sites, colored by root.

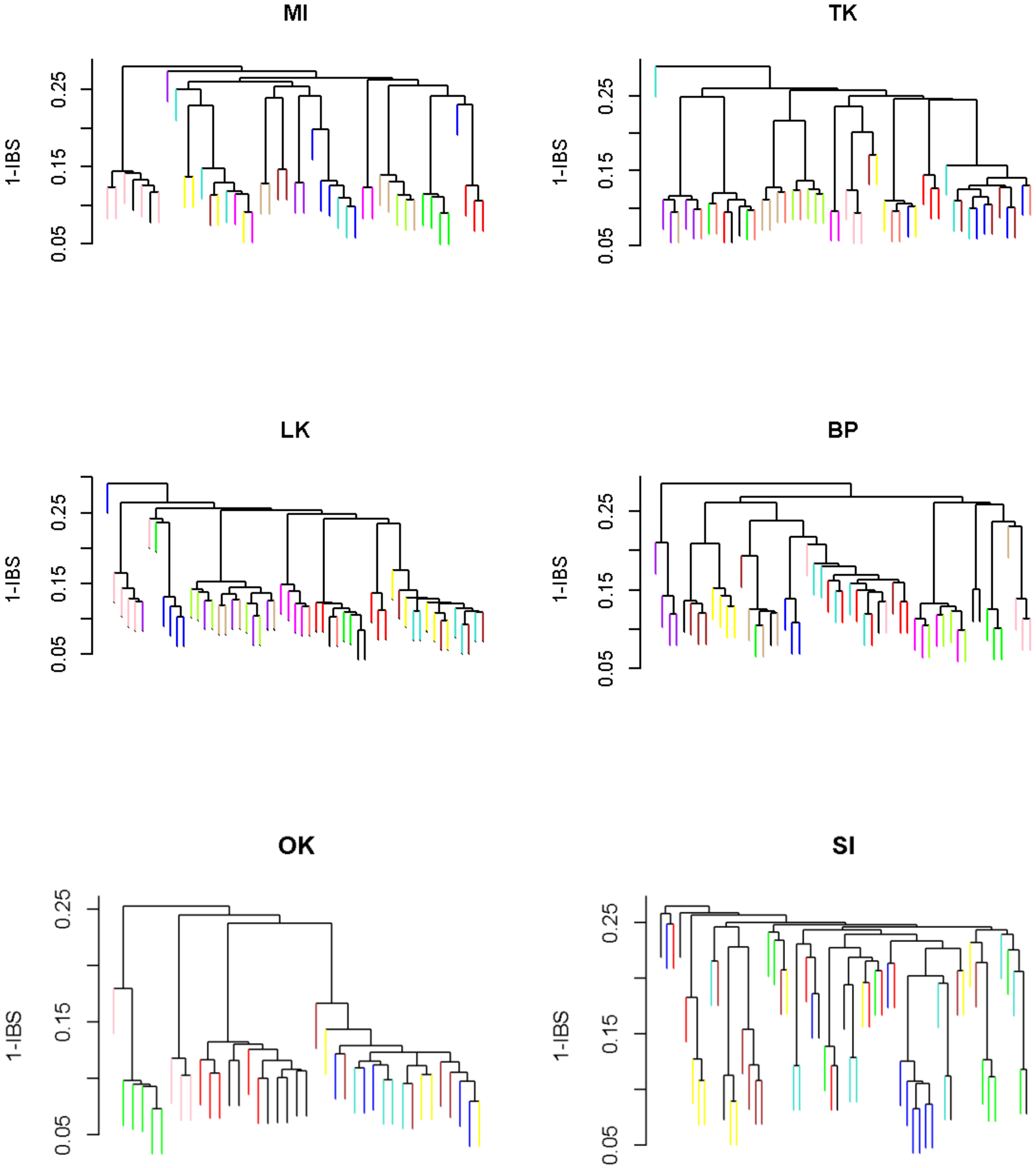

Figure S12.MDS biplot of host prevosti distance for different filtering thresholds.

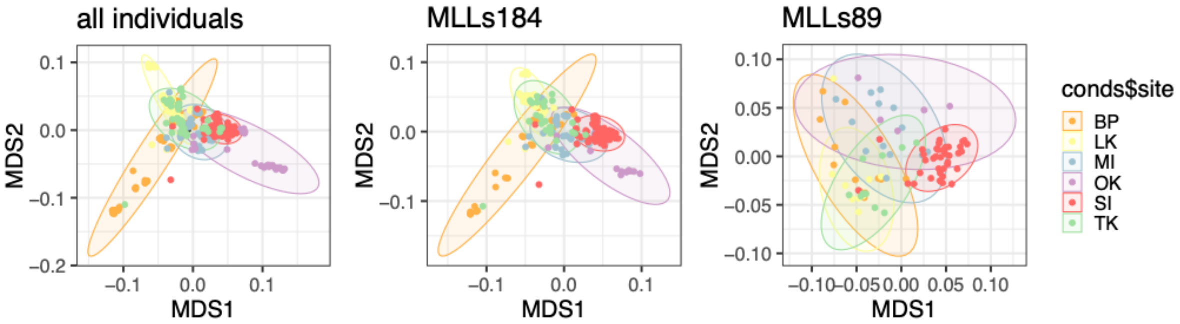


Figure S13.Pairwise Fst (fixation indices) between sites.

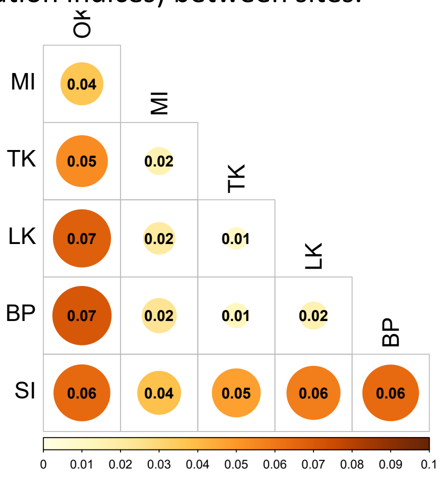


Figure S14.Mean beta diversity between sites.

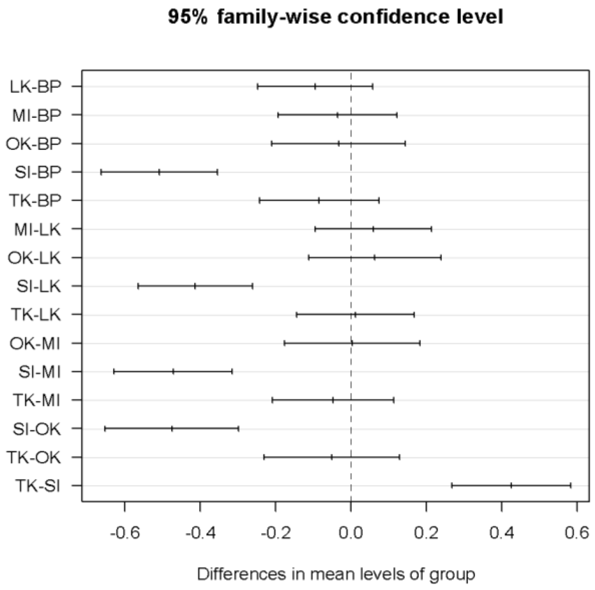

Figure S15. Relative abundance of symbiont profiles within clonal groups. Bolded boxes represent clonal groups (genetic distance <2.3%), with each bar representing the symbiont community composition of clonemates within that group.

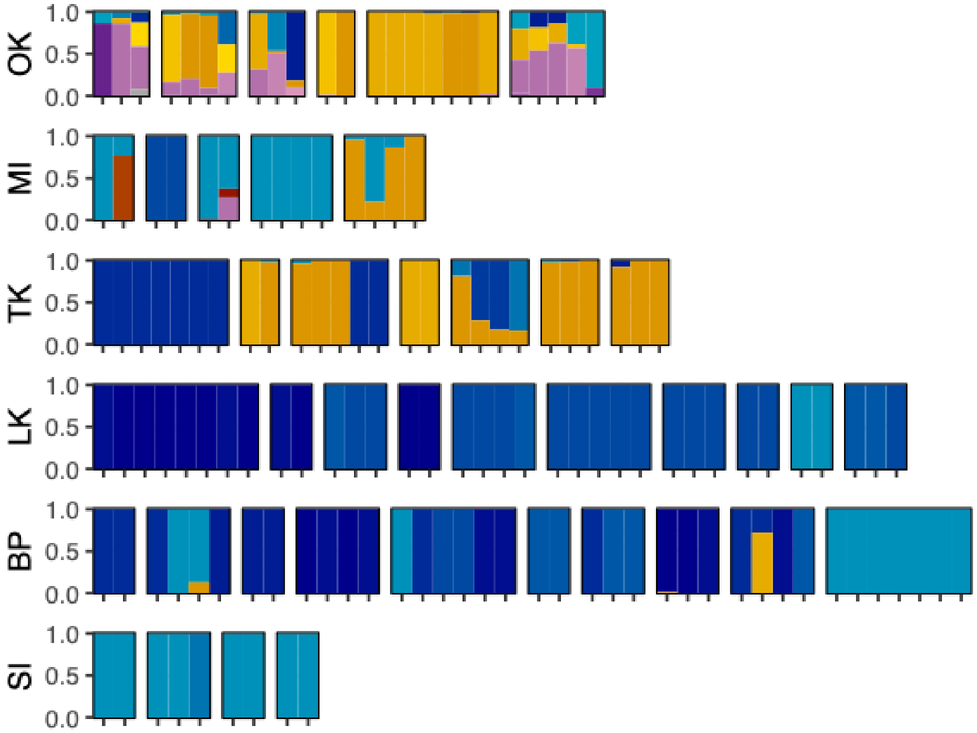


Supplemental tables

Table S1. GPS coordinates of sampling sites.

| Site | Latitude | Longitude |
|---|---|---|
| Stock Island (SI) | 24.565532 | -81.738498 |
| Big Pine (BP) | 24.705185 | -81.35676 |
| Long Key (LK) | 24.824003 | -80.810071 |
| Tavernier Key (TK) | 25.02205 | -80.512955 |
| Miami (MI) | 25.747922 | -80.144946 |
| Otter Key (OK) | 27.313934 | -82.569941 |

Table S2. Pairwise comparisons of residual variance in Fis between sampling locations.

| comp | diff | lwr | upr | p adj |
|---|---|---|---|---|
| MI-OK | -0.0976001 | -0.1298895 | -0.0653108 | 0 |
| TK-OK | -0.0598157 | -0.0923283 | -0.0273031 | 2.41E-06 |
| LK-OK | -0.0826373 | -0.1153654 | -0.0499093 | 6.75E-12 |
| BP-OK | -0.0714382 | -0.1040437 | -0.0388328 | 6.73E-09 |
| SI-OK | -0.1330204 | -0.1652788 | -0.1007619 | 0 |
| TK-MI | 0.03778445 | 0.00743297 | 0.06813593 | 0.00522999 |
| LK-MI | 0.01496282 | -0.0156194 | 0.045545 | 0.73033204 |
| BP-MI | 0.02616192 | -0.004289 | 0.05661286 | 0.13980402 |
| SI-MI | -0.0354202 | -0.0654993 | -0.0053411 | 0.01029545 |
| LK-TK | -0.0228216 | -0.0536394 | 0.00799613 | 0.28157841 |
| BP-TK | -0.0116225 | -0.0423101 | 0.01906501 | 0.88965086 |
| SI-TK | -0.0732047 | -0.1035233 | -0.0428861 | 9.30E-11 |
| BP-LK | 0.01119909 | -0.0197166 | 0.04211482 | 0.90705789 |
| SI-LK | -0.050383 | -0.0809326 | -0.0198335 | 3.88E-05 |
| SI-BP | -0.0615821 | -0.0920003 | -0.0311639 | 1.23E-07 |

Table S3. Clones found across roots.

| MLL (2.3%) | N | rt1 | rt2 | rt3 | rt4 |
|---|---|---|---|---|---|
| MLG.15: | 7 | TK_sub1_rt1 | TK_sub1_rt2 | TK_sub1_rt3 | |
| MLG.34: | 8 | LK_sub3_rt4 | LK_sub3_rt5 | | |
| MLG.45: | 2 | MI_sub1_rt1 | MI_sub3_rt1 | | |
| MLG.61: | 5 | OK_sub2_rt2 | OK_sub2_rt3 | | |
| MLG.66: | 3 | OK_sub2_rt2 | OK_sub2_rt3 | | |
| MLG.81: | 3 | LK_sub2_rt1 | LK_sub2_rt3 | | |
| MLG.94: | 2 | OK_sub1_rt2 | OK_sub1_rt3 | | |
| MLG.101: | 9 | OK_sub1_rt1 | OK_sub1_rt2 | OK_sub1_rt3 | OK_sub1_rt4 |
| MLG.133: | 5 | TK_sub2_rt1 | TK_sub2_rt2 | TK_sub2_rt3 | TK_subNA_rt[illegible] |
| MLG.148: | 4 | BP_sub2_rt1 | BP_sub2_rt3 | BP_sub3_rt4 | |
| MLG.164: | 6 | BP_sub1_rt1 | BP_sub2_rt2 | BP_sub2_rt3 | BP_sub2_rt4 |
| MLG.167: | 4 | TK_sub1_rt2 | TK_sub1_rt4 | TK_subNA_rtNA | |
| MLG.179: | 3 | BP_sub1_rt1 | BP_sub2_rt2 | | |
| MLG.197: | 2 | MI_sub1_rt1 | MI_sub1_rt2 | | |
| MLG.213: | 5 | LK_sub1_rt1 | LK_sub1_rt3 | LK_sub1_rt4 | |
| MLG.215: | 3 | LK_sub1_rt1 | LK_sub1_rt3 | | |
| MLG.216: | 2 | LK_sub1_rt1 | LK_sub1_rt3 | | |
| MLG.231: | 4 | BP_sub1_rt3 | BP_sub2_rt3 | | |
| MLG.255: | 4 | MI_sub2_rt3 | MI_sub2_rt4 | | |
| MLG.264: | 4 | TK_sub3_rt1 | TK_sub3_rt3 | TK_subNA_rtNA | |
| MLG.291: | 7 | BP_sub3_rt1 | BP_sub3_rt3 | | |

Table S4. Clones found across subsites.

| MLL (2.3%) | N | sub1 | sub2 |
|---|---|---|---|
| MLG.45 | 2 | MI_sub1 | MI_sub3 |
| MLG.148 | 4 | BP_sub2 | BP_sub3 |
| MLG.164 | 6 | BP_sub1 | BP_sub2 |
| MLG.179 | 3 | BP_sub1 | BP_sub2 |
| MLG.231 | 4 | BP_sub1 | BP_sub2 |

Table S5. Number of genets present on each mangrove root. Note that SI samples were collected from docks and not mangrove roots so "roots" within this sampling location represents a ~1m$^2$ area. Roots at all other sites are representative of individual mangrove roots.

| root | n_genets |
|---|---|
| SI_sub1_rt1 | 7 |
| SI_sub3_rt1 | 7 |
| SI_sub3_rt2 | 7 |
| SI_sub2_rt2 | 6 |
| SI_sub2_rt1 | 5 |
| BP_sub2_rt3 | 4 |
| SI_sub1_rt2 | 4 |
| BP_sub1_rt3 | 3 |
| BP_sub2_rt4 | 3 |
| MI_sub1_rt1 | 3 |
| BP_sub2_rt1 | 2 |
| BP_sub3_rt2 | 2 |
| BP_sub3_rt4 | 2 |
| LK_sub1_rt2 | 2 |
| LK_sub1_rt4 | 2 |
| LK_sub2_rt1 | 2 |
| LK_sub2_rt2 | 2 |
| LK_sub3_rt1 | 2 |
| LK_sub3_rt3 | 2 |
| MI_sub1_rt2 | 2 |
| MI_sub1_rt3 | 2 |
| MI_sub1_rt4 | 2 |
| MI_sub3_rt1 | 2 |
| MI_sub3_rt2 | 2 |
| MI_sub3_rt4 | 2 |
| OK_sub2_rt4 | 2 |
| TK_sub1_rt1 | 2 |
| TK_sub1_rt2 | 2 |
| TK_sub1_rt4 | 2 |
| TK_sub2_rt2 | 2 |
| TK_sub3_rt3 | 2 |

| root | n_genets |
|---|---|
| BP_sub1_rt1 | 1 |
| BP_sub1_rt2 | 1 |
| BP_sub1_rt4 | 1 |
| BP_sub2_rt2 | 1 |
| BP_sub3_rt1 | 1 |
| BP_sub3_rt3 | 1 |
| LK_sub1_rt1 | 1 |
| LK_sub1_rt3 | 1 |
| LK_sub2_rt3 | 1 |
| LK_sub3_rt2 | 1 |
| LK_sub3_rt4 | 1 |
| LK_sub3_rt5 | 1 |
| MI_sub2_rt1 | 1 |
| MI_sub2_rt2 | 1 |
| MI_sub2_rt3 | 1 |
| MI_sub2_rt4 | 1 |
| MI_sub3_rt3 | 1 |
| OK_sub1_rt1 | 1 |
| OK_sub1_rt2 | 1 |
| OK_sub1_rt3 | 1 |
| OK_sub1_rt4 | 1 |
| OK_sub2_rt1 | 1 |
| OK_sub2_rt2 | 1 |
| OK_sub2_rt3 | 1 |
| TK_sub1_rt3 | 1 |
| TK_sub2_rt1 | 1 |
| TK_sub2_rt3 | 1 |
| TK_sub2_rt4 | 1 |
| TK_sub2_rt5 | 1 |
| TK_sub3_rt1 | 1 |
| TK_sub3_rt2 | 1 |

Table S6. Symbiont clade level PERMANOVA across sampling sites.

| symbiont clade composition | | | | |
|---|---|---|---|---|
| pairs | F.Model | R2 | p.value | p.adjusted |
| MI vs SI | 23.93599616 | 0.184213743 | 0.001 | 0.001 |
| MI vs BP | 18.946153 | 0.152857935 | 0.001 | 0.001 |
| MI vs LK | 26.12735559 | 0.193353562 | 0.001 | 0.001 |
| MI vs TK | 6.821891722 | 0.063270006 | 0.012 | 0.012 |
| MI vs OK | 37.27686765 | 0.307369974 | 0.001 | 0.001 |
| SI vs BP | 1.55218997 | 0.014040337 | 0.259 | 0.259 |
| SI vs LK | 1.054071146 | 0.009241855 | 0.475 | 0.475 |
| SI vs TK | 62.06462294 | 0.371500691 | 0.001 | 0.001 |
| SI vs OK | 229.0346168 | 0.722427788 | 0.001 | 0.001 |
| BP vs LK | 2.558801958 | 0.022336145 | 0.015 | 0.015 |
| BP vs TK | 53.42105975 | 0.339351417 | 0.001 | 0.001 |
| BP vs OK | 192.379017 | 0.688595082 | 0.001 | 0.001 |
| LK vs TK | 66.75763556 | 0.382001252 | 0.001 | 0.001 |
| LK vs OK | 246.0634039 | 0.730021121 | 0.001 | 0.001 |
| TK vs OK | 12.61500629 | 0.131935423 | 0.001 | 0.001 |

Table S7. Symbiont ITS2-type level PERMANOVA across sampling sites.

| ITS2 type composition | | | | |
|---|---|---|---|---|
| pairs | F.Model | R2 | p.value | p.adjusted |
| MI vs SI | 17.788543 | 0.143701045 | 0.001 | 0.001 |
| MI vs BP | 15.28495573 | 0.12707288 | 0.001 | 0.001 |
| MI vs LK | 19.4146462 | 0.151187164 | 0.001 | 0.001 |
| MI vs TK | 19.80513185 | 0.163942802 | 0.001 | 0.001 |
| MI vs OK | 58.05322389 | 0.408672343 | 0.001 | 0.001 |
| SI vs BP | 2.143407483 | 0.019285062 | 0.122 | 0.122 |
| SI vs LK | 1.054071146 | 0.009241855 | 0.493 | 0.493 |
| SI vs TK | 62.93352819 | 0.374752611 | 0.001 | 0.001 |
| SI vs OK | 228.72783 | 0.722158927 | 0.001 | 0.001 |
| BP vs LK | 2.53014844 | 0.022091549 | 0.013 | 0.013 |
| BP vs TK | 53.44833465 | 0.339465862 | 0.001 | 0.001 |
| BP vs OK | 193.7723294 | 0.690140406 | 0.001 | 0.001 |
| LK vs TK | 66.75763556 | 0.382001252 | 0.001 | 0.001 |
| LK vs OK | 243.3735822 | 0.727849313 | 0.001 | 0.001 |
| TK vs OK | 15.52850172 | 0.15760416 | 0.001 | 0.001 |

Table S8. Symbiont ITS2-profile level PERMANOVA across sampling sites.

| ITS2 profile composition | | | | |
|---|---|---|---|---|
| pairs | F.Model | R2 | p.value | p.adjusted |
| MI vs SI | 23.25365223 | 0.179907119 | 0.001 | 0.001 |
| MI vs BP | 6.560741131 | 0.058808691 | 0.001 | 0.001 |
| MI vs LK | 6.33278584 | 0.054908808 | 0.002 | 0.002 |
| MI vs TK | 17.16599972 | 0.145270211 | 0.001 | 0.001 |
| MI vs OK | 12.27820266 | 0.127528374 | 0.001 | 0.001 |
| SI vs BP | 24.15953271 | 0.181432994 | 0.001 | 0.001 |
| SI vs LK | 52.95479802 | 0.319091696 | 0.001 | 0.001 |
| SI vs TK | 64.51926852 | 0.380601386 | 0.001 | 0.001 |
| SI vs OK | 49.10507869 | 0.358156526 | 0.001 | 0.001 |
| BP vs LK | 9.757188328 | 0.080136446 | 0.001 | 0.001 |
| BP vs TK | 11.8593688 | 0.102360033 | 0.001 | 0.001 |
| BP vs OK | 9.998706273 | 0.103080821 | 0.001 | 0.001 |
| LK vs TK | 21.97204918 | 0.16905211 | 0.001 | 0.001 |
| LK vs OK | 15.18439254 | 0.14300023 | 0.001 | 0.001 |
| TK vs OK | 8.719682113 | 0.095068822 | 0.001 | 0.001 |

Table S9. Clade level differences in beta diversity across sites.

| ITS2 clade beta diversity | | | | |
|---|---|---|---|---|
| | diff | lwr | upr | p adj |
| LK-BP | -0.0234 | -0.1400 | 0.0932 | 0.9926 |
| **MI-BP** | **0.2549** | **0.1346** | **0.3753** | **0.0000** |
| **OK-BP** | **0.2761** | **0.1403** | **0.4118** | **0.0000** |
| SI-BP | -0.0195 | -0.1376 | 0.0987 | 0.9970 |
| **TK-BP** | **0.4261** | **0.3052** | **0.5471** | **0.0000** |
| **MI-LK** | **0.2783** | **0.1600** | **0.3967** | **0.0000** |
| **OK-LK** | **0.2995** | **0.1655** | **0.4334** | **0.0000** |
| SI-LK | 0.0039 | -0.1122 | 0.1200 | 1.0000 |
| **TK-LK** | **0.4495** | **0.3306** | **0.5685** | **0.0000** |
| OK-MI | 0.0212 | -0.1161 | 0.1584 | 0.9979 |
| **SI-MI** | **-0.2744** | **-0.3942** | **-0.1545** | **0.0000** |
| **TK-MI** | **0.1712** | **0.0486** | **0.2938** | **0.0011** |
| **SI-OK** | **-0.2955** | **-0.4308** | **-0.1603** | **0.0000** |
| **TK-OK** | **0.1501** | **0.0123** | **0.2878** | **0.0238** |
| **TK-SI** | **0.4456** | **0.3252** | **0.5660** | **0.0000** |

Table S10. ITS2 type level differences in beta diversity across sites.

| ITS2 type beta diversity | | | | |
|---|---|---|---|---|
| **Pairs** | **diff** | **lwr** | **upr** | **p adj** |
| LK-BP | -0.0234 | -0.1418 | 0.0950 | 0.9931 |
| **MI-BP** | **0.2549** | **0.1327** | **0.3771** | **0.0000** |
| **OK-BP** | **0.2835** | **0.1457** | **0.4213** | **0.0000** |
| SI-BP | -0.0195 | -0.1394 | 0.1005 | 0.9973 |
| **TK-BP** | **0.4229** | **0.3001** | **0.5457** | **0.0000** |
| **MI-LK** | **0.2783** | **0.1581** | **0.3985** | **0.0000** |
| **OK-LK** | **0.3069** | **0.1709** | **0.4429** | **0.0000** |
| SI-LK | 0.0039 | -0.1139 | 0.1218 | 1.0000 |
| **TK-LK** | **0.4463** | **0.3255** | **0.5671** | **0.0000** |
| OK-MI | 0.0286 | -0.1108 | 0.1679 | 0.9918 |
| **SI-MI** | **-0.2744** | **-0.3961** | **-0.1527** | **0.0000** |
| **TK-MI** | **0.1680** | **0.0435** | **0.2925** | **0.0018** |
| **SI-OK** | **-0.3030** | **-0.4403** | **-0.1656** | **0.0000** |
| TK-OK | 0.1394 | -0.0005 | 0.2793 | 0.0514 |
| **TK-SI** | **0.4423** | **0.3201** | **0.5646** | **0.0000** |

Table S11. ITS2 profile level differences in beta diversity across sites.

| ITS2 profile beta diversity | | | | |
|---|---|---|---|---|
| **Pairs** | **diff** | **lwr** | **upr** | **p adj** |
| LK-BP | -0.0957 | -0.2484 | 0.0571 | 0.4696 |
| MI-BP | -0.0365 | -0.1941 | 0.1211 | 0.9857 |
| OK-BP | -0.0332 | -0.2109 | 0.1446 | 0.9947 |
| **SI-BP** | **-0.5091** | **-0.6638** | **-0.3544** | **0.0000** |
| TK-BP | -0.0843 | -0.2427 | 0.0741 | 0.6474 |
| MI-LK | 0.0592 | -0.0958 | 0.2141 | 0.8832 |
| OK-LK | 0.0625 | -0.1129 | 0.2379 | 0.9104 |
| **SI-LK** | **-0.4135** | **-0.5655** | **-0.2614** | **0.0000** |
| TK-LK | 0.0113 | -0.1444 | 0.1671 | 0.9999 |
| OK-MI | 0.0033 | -0.1764 | 0.1830 | 1.0000 |
| **SI-MI** | **-0.4726** | **-0.6295** | **-0.3157** | **0.0000** |
| TK-MI | -0.0478 | -0.2084 | 0.1128 | 0.9569 |
| **SI-OK** | **-0.4760** | **-0.6531** | **-0.2988** | **0.0000** |
| TK-OK | -0.0512 | -0.2316 | 0.1292 | 0.9650 |
| **TK-SI** | **0.4248** | **0.2671** | **0.5825** | **0.0000** |

Table S12. Spatial correlation (Moran's I) of ITS2 symbiont profiles including all individuals.

| **All individuals** | | | | | |
|---|---|---|---|---|---|
| **ITS2 profile** | **p.val** | **I.obs** | **I.exp** | **I.sd** | **p.adj** |
| A4 | 0 | 0.598477269 | -0.003424658 | 0.029748237 | 0 |
| A4-A4ct-A4cs-A4q | 0 | 0.529477808 | -0.003424658 | 0.029632945 | 0 |
| A4/A4m | 0 | 0.432133709 | -0.003424658 | 0.029379224 | 0 |
| A4-A4ct-A4cs-A1il | 0 | 0.280813894 | -0.003424658 | 0.029168747 | 0 |
| A4-A4q-A4cw | 0 | 0.312662673 | -0.003424658 | 0.029192868 | 0 |
| A4-A4q-A1il | 0.000914849 | 0.092272419 | -0.003424658 | 0.02886369 | 0.000914849 |
| B2 | 0 | 0.584841433 | -0.003424658 | 0.029449059 | 0 |
| B2-B2al | 0 | 0.545675375 | -0.003424658 | 0.028986965 | 0 |

Table S13. Spatial correlation (Moran's I) of ITS2 symbiont profiles after removing clones.

| **No clones** | | | | | |
|---|---|---|---|---|---|
| **ITS2 profile** | **p.val** | **I.obs** | **I.exp** | **I.sd** | **p.adj** |
| A4 | 1.39E-10 | 0.560954047 | -0.013333333 | 0.089489203 | 8.32E-10 |
| A4-A4ct-A4cs-A4q | 9.06E-05 | 0.326091217 | -0.013333333 | 0.086708064 | 0.000181133 |
| A4/A4m | 0.896527177 | -0.023632418 | -0.013333333 | 0.079193508 | 0.896527177 |
| A4-A4q-A1il | 0.573006587 | -0.059383772 | -0.013333333 | 0.081703445 | 0.687607905 |
| B2 | 2.44E-06 | 0.37391793 | -0.013333333 | 0.082163916 | 7.32E-06 |
| B2-B2al | 0.273055144 | 0.072245657 | -0.013333333 | 0.078079173 | 0.409582716 |

Table S14. Summary of PACo model results.

| **Genera** | **null model** | **m2xy** | **p** |
|---|---|---|---|
| **Symbiodinium** | **r0** | **5.323619** | **0** |
| Symbiodinium | c0 | 5.323619 | 0.425 |
| **Symbiodinium** | **swap** | **5.323619** | **0** |
| **Symbiodinium** | **backtrack** | **5.323619** | **0** |
| **Breviolum** | **r0** | **1.793123** | **0.002** |
| Breviolum | c0 | 1.793123 | 0.743 |
| **Breviolum** | **swap** | **1.793123** | **0.004** |
| **Breviolum** | **backtrack** | **1.793123** | **0** |
| Cladocopium | r0 | 0.6915839 | 0.884 |
| Cladocopium | c0 | 0.6915839 | 0.57 |
| Cladocopium | swap | 0.6915839 | 0.918 |
| Cladocopium | backtrack | 0.6915839 | 0.897 |
